# Astrocyte stimulation reopens the window of the critical period for the experience-dependent plasticity of retinogeniculate synapses

**DOI:** 10.64898/2026.01.14.699178

**Authors:** Congyun Jin, Yasuhiro Go, Junichi Nabekura, Madoka Narushima

## Abstract

A specific time window during which neural circuits undergo dramatic changes in response to individual experiences is known as the critical period (CP). Developing methods to reopen the CP after its closure could lead to the establishment of effective treatment approaches. The retinogeniculate (RG) synapses in the dorsal lateral geniculate nucleus (dLGN) undergo experience-dependent remodeling only during the limited period of development. We investigated the effects of astrocyte manipulation on the restoration of RG synapse plasticity after the CP termination. Astrocytic stimulation with the designer receptors exclusively activated by designer drugs (DREADD) after the CP closure caused RG synapse remodeling in the dark-reared mice but not in those reared in normal conditions. Single-nucleus RNA sequencing (snRNA-seq) of dLGN astrocytes implicated the underlying mechanisms of astrocyte-induced RG synaptic plasticity. Therefore, astrocytes can regulate the susceptibility of RG synapses to experience-dependent plasticity after CP termination.

## Introduction

Central synapses are reorganized in response to the individual’s experience during the limited developmental period known as the critical period (CP) (Hooks and Chen, 2020; Reh et al., 2020), a phenomenon widely observed across brain areas. In addition, several neurological conditions are known to exhibit time windows of heightened therapeutic sensitivity, reminiscent of developmental critical periods (Zeiler, 2019). Therefore, controlling the plasticity of neural circuits after CP termination and guiding them in the right direction is a major goal of neuroscience research. The CP of sensory neural circuits, in which the onset and offset of plasticity are strictly controlled, is a suitable model for developing methods to manipulate the CP.

The plasticity of neural circuits is strongly influenced by glial cells (Stogsdill and Eroglu, 2017). The astrocyte is a type of glial cell that plays a crucial role in maintaining homeostasis within the central nervous system (Perez-Catalan et al., 2021). Recent studies revealed that astrocytes contribute to the termination of the CP by promoting stabilization of the extracellular matrix (ECM) (Ribot et al., 2021) or by stabilizing dendritic morphology via neuroligin-neurexin signaling (Ackerman et al., 2021). In contrast, astrocytes can also promote neuronal plasticity even in the adult brain (Kim et al., 2016). We recently reported that in the chronic period of neuropathic pain, artificial stimulation of astrocytes by a Gq-activated type of designer receptors exclusively activated by designer drugs (DREADD), hM3D (Armbruster et al., 2007), in combination with suppression of peripheral nerve activity, enhances pruning of newly formed synapses and then relieves allodynia (Takeda et al., 2022). Thus, astrocytes can regulate neuronal network organization in both stabilizing and remodeling directions. In general, experience-dependent plasticity is a multidirectional phenomenon, with synapse strengthening or weakening depending on the synapse’s role, and can be reversed when sensory experiences are regained during the CP. However, it is not straightforward to independently analyze the processes underlying each of these plastic changes during the CP. Therefore, whether astrocytes can reinstate multidirectional synaptic plasticity, encompassing both synaptic strengthening and weakening, has not been investigated at the synaptic level. In this study, we aimed to determine, at the synaptic level, whether stimulating astrocytes can restore multidirectional neuronal circuit plasticity after CP closure and to elucidate the underlying mechanisms. To this end, we focused on the synapses between retinal ganglion cells (RGCs) and thalamocortical (TC) neurons in the dorsal lateral geniculate nucleus (dLGN), a system with a well-defined CP that allows examination of bidirectional circuit plasticity at the synaptic level.

In mice, the retinogeniculate (RG) synapses can be bidirectionally reorganized in response to their visual experience during a strictly limited CP which starts ∼postnatal days of (P) 20 and closes by P30 (Hooks and Chen, 2006, 2008). When visual experience is blocked by dark rearing (DR), existing RG synapses are weakened, and new synapses are formed excessively (Hooks and Chen, 2006, 2008). In contrast, exposure to light after DR can reverse the plasticity and promote synapse pruning (the recovery period) (Hooks and Chen, 2008). According to those features, the experience-dependent RG synapse remodeling is referred to as a type of homeostatic plasticity (Wen and Turrigiano, 2024), and an excellent model for studying the regulatory mechanisms of bidirectional plasticity, which has a clear CP. We found that combining astrocyte stimulation with visual deprivation can induce RG synapse remodeling after the CP termination, thereby reopening the CP window for experience-dependent plasticity. Interestingly, synapse pruning after the remodeling was promoted only when astrocytes continued to be activated, and mice regained visual experience. In addition, we observed changes in astrocyte gene expression following DREADD stimulation, which may underlie reopening of the CP. Our results clearly showed that artificial stimulation of astrocytes can reopen the CP for RG synapse remodeling and suggest possible molecular mechanisms underlying the astrocytic control of bidirectional plasticity.

## Results

### 1. The critical period of experience-dependent plasticity in RG synapses

In this study, we utilized Chx10-cre; ChR2 mice to selectively activate RGC axons innervating the dLGN excitatory TC neurons (Litvina and Chen, 2017). We observed reorganization of RG synaptic connectivity after DR from P21 in Chx10-cre; ChR2 mice (Figures 1A and 1B) (Table S1). When the intensity of stimulus by blue LED light gradually increased during patch clamp recording from TC neurons, the amplitude of retinal excitatory synaptic currents (EPSCs) increased in a stepwise manner. For the sake of the high release probability of RGC axon terminals, the RG-EPSC is generated in an all-or-none fashion. Therefore, the single-step amplitude can be regarded as synaptic transmission originating from a single RGC axon (Chen and Regehr, 2000). On the postnatal days of (P) 21, the mice Chx10-cre; ChR2 were replaced in a box with 24-hr darkness (dark-reared/DR) or 12-hr light/dark cycle (light-reared/LR) for 8 to 15 days. The distribution of the step number of RG-EPSCs in the DR group (“CP DR”, n = 27 cells) shifted toward increasing compared to the age-matched control reared in a normal cycle (“CP LR”, n = 20 cells, p = 0.002) (Figure 1C). The single fiber (SF) amplitude of RG-EPSCs, that is, the amplitude of each step of RG-EPSCs, decreased after DR (109 responses, p < 0.001) (Figure 1G). We calculated the SF fraction (SFF), which is the ratio between an SF amplitude and the maximum amplitude of AMPR-EPSCs recorded from the same cell, to evaluate the relative strength of an SF input compared with the maximum amplitude in a single TC neuron. The maximum amplitude was not affected (p = 0.188) (Figure 1H), and the SFF was reduced by DR (P <0.001) (Figure 1I), suggesting an increase in innervating RGC axons and a weakening of each RG synapse. Next, we checked whether it was possible to reproduce the CP of RG synapse remodeling in Chx10-cre; ChR2 mice. DR initiated from P28 (“Post-CP DR”) did not cause reorganization of RG synapses (Figures 1D and 1E). Neither the RG-EPSC step number (n = 16, p = 0.47) (Figure 1F) nor the SFF (p = 0.084) (Figure 1I) differed from that of the LR group (“Post-CP LR”). These results confirm a narrow window of the CP for experience-dependent RG synapse remodeling.

**Figure 1.**
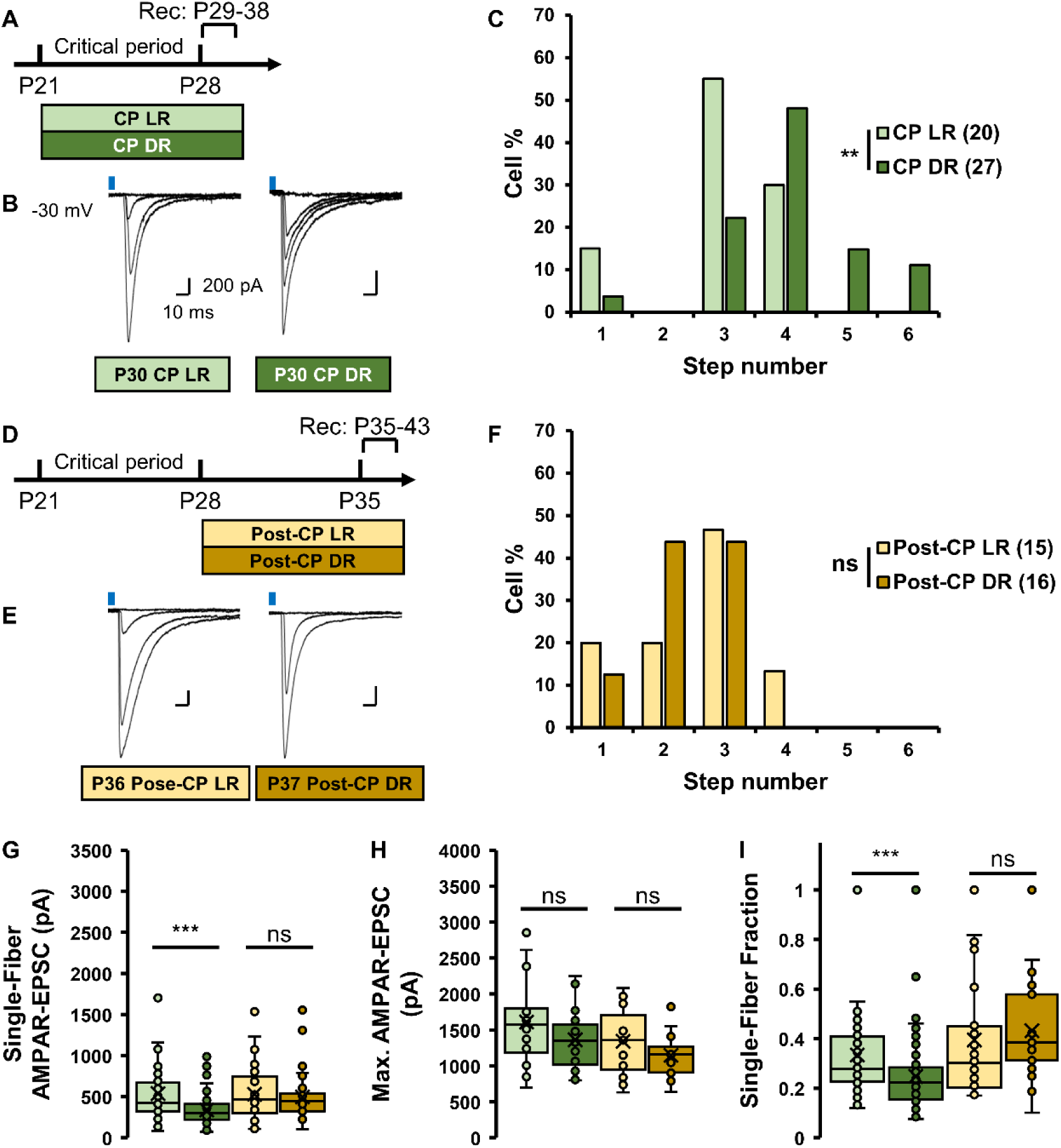
Experience-dependent remodeling of retinogeniculate synapses during the critical period with ChR2 stimulation of retinal fibers. **(A**–**F)** Timeline (A and D), representative traces of RG-EPSCs (B and E) and histograms showing the number of RG-EPSC steps (C and F) in Chx10-Cre; ChR2 mice subjected to visual deprivation during the CP (A - C) and post–CP (D - F). After one week of DR or LR, AMPAR-mediated RG-EPSCs were recorded from TC neurons in the dLGN in response to LED blue light stimulation at a holding potential of –30 mV. Calibration bars: 200 pA, 10 ms. Numbers in parentheses in C and F indicate the number of cells analyzed per group. **(G**–**I)** Properties of RG-EPSCs in Chx10-Cre;ChR2 mice. Single-fiber EPSC amplitude (G) and maximal AMPAR-mediated EPSC amplitude (H) were recorded at a holding potential of –30 mV. ***p < 0.001; **p < 0.01; *p < 0.05 (Mann–Whitney U-test); ns, not significant.

### 2. DREADD-stimulation of astrocytes after the closure of CP resulted in the regaining of RG synapse plasticity

The disappearance of experience-dependent RG synapse remodeling suggests that the susceptibility to synaptic plasticity changes after P28. Therefore, we stimulated astrocytes, which can control neuronal plasticity (Perez-Catalan et al., 2021), as potential regulators for RG synapse remodeling. To artificially stimulate astrocytes in the dLGN, we developed an AAV carrying hM3D and mScarlet genes or mScarlet gene alone that are translated under the control of the human ALDH1L1 promoter (Koh et al., 2017) (Figure S1). We confirmed that two weeks after injection of the AAV, mScarlet was dominantly expressed in S100β-positive astrocytes in the dLGN (Figures S1B, and S1C). We also confirmed that the application of the artificial ligand, CNO, can induce Ca2+ elevation in the hippocampal culture slices (Figures S1D and S1E). The either of AAVs was injected into the dLGN at P14, then two weeks later, from P28, mice were reared under DR or LR conditions, with/without astrocyte stimulation (hM3D and Control AAV; Clozapine N-oxide (CNO) and saline), until RG synapse remodeling was quantified by *in vitro* electrophysiology (Figures 2A and 2B) (Table S1).

**Figure 2.**
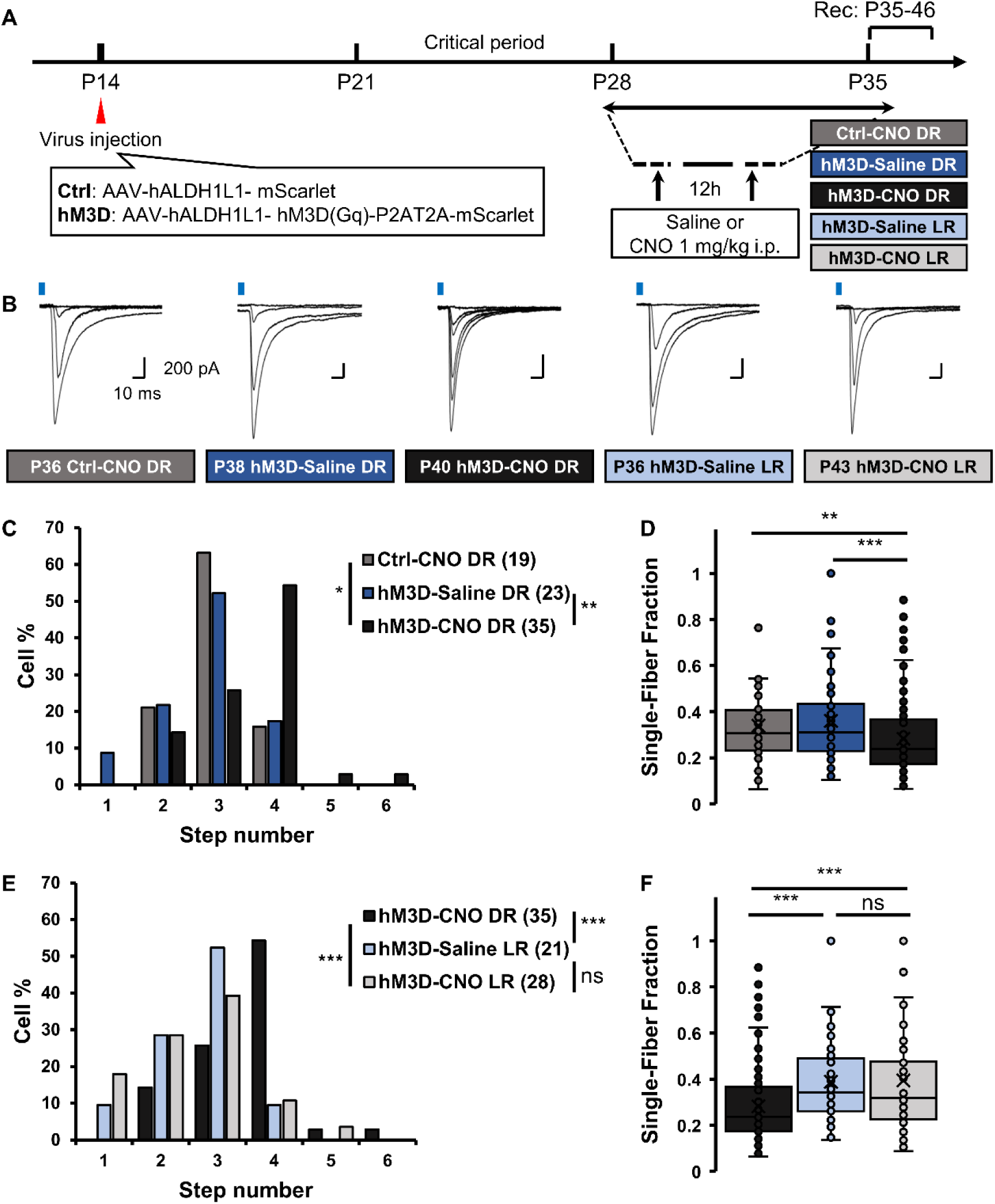
Chemogenetic activation of astrocytes in the dLGN restores synaptic reorganization after the critical period. **(A)** Timeline of the DREADD-based manipulation of astrocyte activity. AAV viruses were injected into the dLGN at P14. Two weeks later, mice were subjected to 1 week of DR or LR. CNO (1 mg/kg) was administered twice daily during this period. RG-EPSCs were then recorded from TC neurons in the dLGN. **(B)** Representative traces of RG-EPSCs recorded at a holding potential of –30 mV after indicated manipulations. Calibration bars: 200 pA, 10 ms. **(C and D)** Quantification of RG-EPSC step numbers (C) and the single-fiber fraction (D) non-indicated conditions. **(E and F)** The same analysis was performed with (C) and (D) in indicated conditions. Numbers in parentheses in (C) and (E) indicate the number of cells analyzed per group. ***p < 0.001; **p < 0.01; *p < 0.05 (Mann–Whitney U-test); ns, not significant.

RG-EPSCs recorded from TC neurons in the dLGN expressing hM3D in astrocytes but treated with saline during DR (“hM3D-Saline DR”, n = 23) or expressing a control vector with CNO treatment (“hM3D-CNO DR”, n = 19) showed no clear evidence of synapse remodeling in response to DR (Figures 2C and 2D). The distribution of step number, maximum amplitude, or SFF was not different from that of the LR-saline mice (Figures 2E and 2F, Figures S2C and S2D). Interestingly, RG-EPSCs of hM3D-expressing mice with DR and CNO treatment (“hM3D-CNO DR”, n = 35) showed clearly a larger number of steps than those in Ctrl-CNO DR (p = 0.013) or hM3D-Saline DR (p = 0.004) (Figure 2C). The maximum amplitude was not different among the conditions (Figure S2B), but SF amplitude decreased in the hM3D-CNO DR group (p = 0.038 compared with the Ctrl-CNO DR group) (Figure S2A). As a result, the SFF was significantly decreased in hM3D-CNO DR mice but not in hM3D-Saline DR (p = 0.001) or in Ctrl-CNO DR (p = 0.005) (Figure 2D). Together with the increase in the step number, the reduction of the SFF suggests a weakening of the synaptic strength in each retinal fiber and an increment in the number of innervating RGC axons to a single TC neuron, as occurred by DR during the CP.

Importantly, the step number distribution or the SFF of hM3D-expressing mice with LR and CNO treatment (“hM3D-CNO LR”, n = 28) was not different from the LR-saline group (“hM3D-Saline LR”, n = 21) (p = 0.702 for the step number and 0.489 for the single fiber fraction) but was significantly different from the hM3D-CNO DR group (p < 0.001 for the step number and p = 0.001 for the SFF) (Figures 2E and 2F), indicating that stimulation of astrocytes alone could not affect the plasticity of RG synapses. These results revealed that astrocyte stimulation enhances RG synapse remodeling in conjunction with changes in sensory experience, even after CP closure.

### 3. Astrocyte stimulation restarted RG synapse pruning with light exposure after the CP termination

Next, we examined whether a “recovery period”—that is, pruning of the re-formed synapses—also occurs after RG synaptic remodeling was reopened by astrocytic stimulation. The cages for the hM3D-CNO DR mice, in which RG synapse remodeling was likely occurring, were returned to the shelf under normal light/dark conditions for at least three days to recover from visual deprivation with saline injection (“Recovery Saline”, n = 27) (Figures 3A and 3B) (Table S1). Although the SF AMPAR-EPSC amplitude was increased (Figure S2E), neither the step number distribution (p = 0.882) (Figure 3C), the maximum amplitude (p = 0.092) (Figure S2F), nor the SFF (p = 0.733) (Figure 3D) was different from that of the hM3D-CNO DR treatment group. This means that reorganized RG synapses failed to be rewired during the recovery period. Interestingly, when CNO was continued to be applied during the recovery period to activate astrocytes in hM3D-expressing mice (“Recovery CNO”, n = 18), the step number of RG EPSCs significantly decreased (p = 0.001) (Figure 3C). Furthermore, the single fiber amplitude increased (p = 0.007) (Figure S2E), but the maximum amplitude was not different with hM3D-CNO DR (p = 0.11) (Figure S2F), resulting in an increase of the SFF (p < 0.001) (Figure 3D) recovery period. These findings suggest the pruning of excess synapses and strengthening of remaining synapses and indicate that bidirectional experience-dependent synapse remodeling, which is a hallmark of experience-dependent RG synapse plasticity, was regained after the CP closure with astrocyte stimulation.

**Figure 3.**
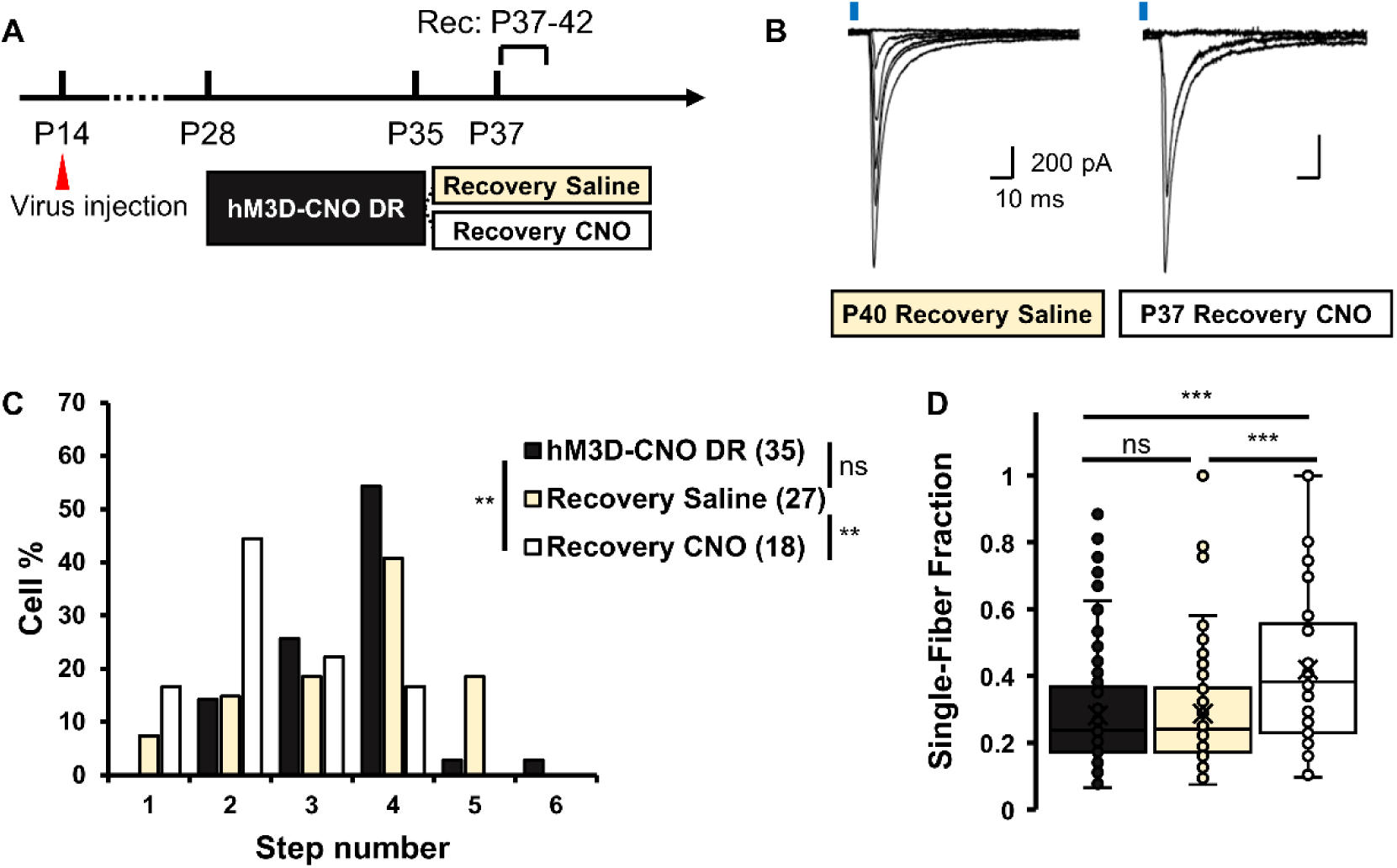
Light exposure-induced recovery from DR–induced synaptic remodeling requires astrocyte activation. **(A)** Timeline for the recovery test. After one week of DR and chemogenetic astrocyte stimulation, mice were returned to a 12-hour light/dark cycle for at least 3 days, with continuous administration of CNO (Recovery CNO group) or saline (Recovery Saline group). **(B)** Representative traces of RG-EPSCs recorded at a holding potential of –30 mV. Calibration bars: 200 pA, 10 ms. **(C and D)** Quantification of RG-EPSC step numbers (C) and the single-fiber fraction (D) in indicated conditions. ***p < 0.001; **p < 0.01; *p < 0.05 (Mann–Whitney U-test); ns, not significant.

### 4. Visual experience-dependent and DREADD stimulation-induced changes in gene expression

Finally, we aimed to clarify the molecular mechanisms by which astrocytic stimulation enhances plasticity by comparing gene expression in astrocytes during the CP and under hM3D activation. We performed single-nucleus RNA sequencing (snRNA-seq) on samples collected from CP, hM3D, or Ctrl-dLGN after DR or LR (Figure 4A). We classified dLGN astrocytes and other cells depending on the expression of marker genes (Figures S3A and S3B) (Kalish et al., 2018).

**Figure 4.**
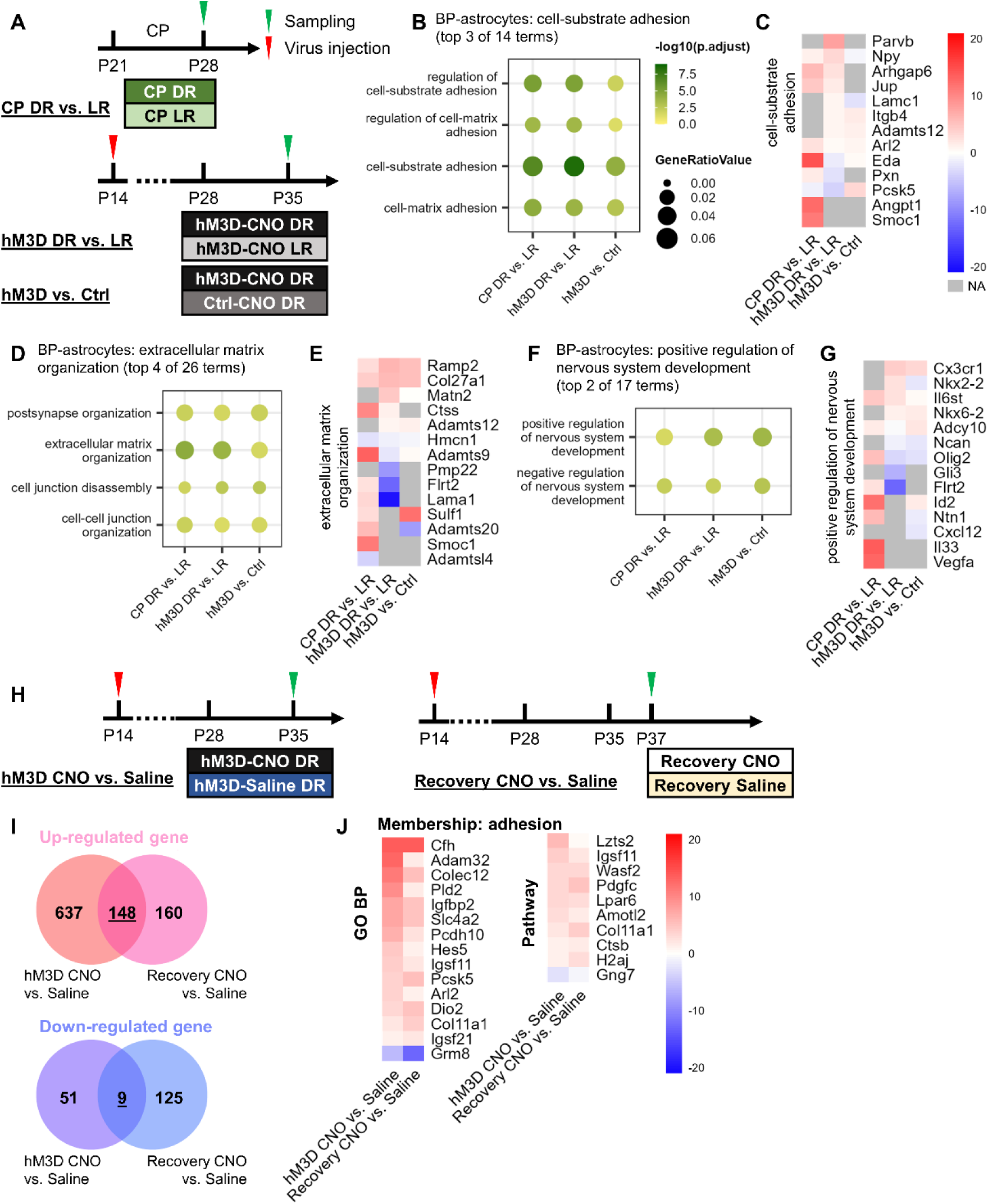
Astrocyte gene expression changes in response to visual experience and chemogenetic activation. **(A)** Timeline of comparisons for snRNA-seq: DR versus LR during the CP (CP DR vs. LR), DR versus LR with astrocyte stimulation after the CP (hM3D DR vs. LR), and astrocyte-stimulated versus unstimulated with DR after the CP (hM3D vs. Ctrl). **(B, D, and F)** Enrichment of clustered GO BP terms identified by semantic similarity analysis (see Figure S4) of ORA. Each cluster groups terms associated with cell-substrate adhesion (B), extracellular matrix organization (C), regulation of nervous system development (E). The top 2–4 terms in each cluster are shown. Circle color represents –log10 (adjusted p-value), and circle size corresponds to the gene ratio associated with each term. **(C, E, and G)** Top 5 genes with high z-score (abs(z_FC) + z_p-value). The colors represent log₂ fold changes, scaled for each gene, and centered at 0. Genes not appearing in the DEG are shown in gray. **(H - J)** Timeline of comparisons for astrocytic genetic alternation by hM3D stimulation irrespective of visual experience. The DEGs of astrocyte-stimulated versus unstimulated during the post-CP with DR (hM3D CNO vs. Saline) and during the recovery period (Recovery CNO vs. Saline) (H) were compared by using the Venn diagram (I) showing the number of overlapping DEGs between hM3D CNO vs. Saline and Recovery CNO vs. Saline. Heatmaps show selected genes mapped to BP terms (J, left) and pathways (J, right) that include “adhesion” in their term name or description. Heatmap colors represent log_2_ fold change, scaled per gene, and centered at 0.

We first hypothesized that stimulation of astrocytes after the CP promoted synapse remodeling by altering gene expression to a state resembling the CP. To verify this possibility, over-representation analysis (ORA) was performed to determine which Gene Ontology (GO) biological process (BP) terms were enriched among CP DR vs. LR, hM3D DR vs. LR, and hM3D vs. Ctrl differently expressed genes (DEGs) in astrocytes (Figure S3C). Focusing on the effects of astrocytes on other cells, we examined parent terms (Figure S4) that represent astrocytic influence on the extracellular environment—namely, the cell-substrate adhesion (GO: 0031589), the extracellular matrix organization (GO: 0030198), and the positive regulation of nervous system development (GO: 0051962) (Figures 4B, D, and F). In the CP DR vs. LR DEGs, we found a clear upregulation of secreted factors that modulate the extracellular environment and cell adhesion, such as *Eda*, *Angpt1*, or *Smoc1*; ECM components, such as *Col27a1* or *Lama1*; enzymes related to ECM remodeling, such as *Ctss*, *Adamts9*, or *Adamts20* (Figures 4C and 4E). In addition, DEGs enriched for positive regulation of nervous system development included cytokines and axon guidance factors such as *Il33*, *Ntn1*, and *Vegfa* (Figure 4G). Thus, the CP-DR astrocytes would affect RG synapse plasticity through remodeling of the ECM and/or cytokine signaling.

As expected, the DEGs of hM3D DR vs. LR, which represented experience-dependent alternation in gene expression after the CP, were mostly different from the CP DR vs. LR. We found suppression of ECM proteins, alterations in metalloproteinase members, and loss of cytokines, all of which were upregulated in CP DR vs. LR DEGs (Figures 4C, E, and G). If DREADD stimulation rejuvenates astrocytes, it compensates for differences in gene expression during and after the CP. However, hM3D stimulation did not compensate for the loss of experience-dependent upregulation of ECM proteins and cytokines. In hM3D vs. Ctrl DEGs, expression of some of the extracellular proteins, such as *Sulf1*, was upregulated, suggesting modification of extracellular signaling through other molecular mechanisms. These results suggest that stimulation of astrocytes after the CP did not reproduce the molecular state of astrocytes during the CP.

The other possibility is that stimulated astrocytes alter the susceptibility of RG synapses to experience-dependent plasticity, making them more likely to reorganize in response to visual experience. If so, regulatory genes for synaptic plasticity in DREADD-stimulated astrocytes changed their expression irrespective of deprivation or recovery of vision, and the direction of synapse remodeling. To find those genes, we compared DEGs of the hM3D CNO vs. saline DR comparison (formation/weakening state) with the Recovery CNO vs. saline comparison (pruning state) (Figure 4H). In astrocytes, we found 148 upregulated and 9 downregulated common genes (Figure 4I), which were functionally diverse. Among these, genes reported to be involved in astrocyte regulation of synaptic transmission or neuronal function, such as *Igfbp2*, *Igsf11*, and *Dio2*, were found in a BP term related to cell adhesion (Figure 4J).

Taken together, astrocyte stimulation does not simply promote astrocyte rejuvenation by returning the molecular state to the CP. Rather, DREADD-stimulated astrocytes likely influence the ECM and neurons through pathways distinct from those engaged during the CP, and these effects may enhance neural circuit plasticity and thereby reopen the CP for bidirectional experience-dependent RG synapse remodeling.

## Discussion

In this study, we successfully reopened the CP of bidirectional experience-dependent RG synapse remodeling through astrocytic stimulation. Astrocytic stimulation not only induced RG synapse reformation and weakening following visual deprivation but also reinstated synapse pruning upon restoration of visual experience. However, astrocytic stimulation alone was insufficient to enhance plasticity; sensory experience manipulation was always required in combination. In addition, our snRNA-seq revealed that astrocytic stimulation did not revert astrocytes to a CP–like state, but it induced changes in the expression of factors that could influence the ECM and neurons.

Because of the presence of a short CP with a clear beginning and end, the RG synapse has the potential to be an important model system for studying how the brain acquires plasticity and how the timing and direction of plasticity are controlled (Liang and Chen, 2020). RG synapses were first reported as a model system for developmental synapse pruning (Chen and Regehr, 2000). Later, its activity-dependent maturation and experience-dependent remodeling toward an immature phenotype of synaptic connectivity was also reported (Hooks and Chen, 2006, 2008). The opening of the CP of RG synapse remodeling is around P20, requiring a few days of visual experience (Hooks and Chen, 2008). RG synapse plasticity is lost by P30 in parallel with the development of activity-dependent maintenance mechanisms (Cheadle et al., 2020; Cheadle et al., 2018; Narushima et al., 2016; Thompson et al., 2016; Tzeng et al., 2023), and then the synapse becomes resistant to visual deprivation (Hooks and Chen, 2008) (Figure 1). In this study, we clearly showed that DREADD stimulation of astrocytes overcame the maintenance mechanisms and reopened the CP for RG synapse remodeling (Figures 2 and 3). There are possibilities that (1) hM3D-stimulated astrocytes switch between functions such as synapse formation and synapse removal in a sensory experience-dependent manner or (2) the effect of astrocytic stimulation on RG synapse plasticity is indirect—regulating the susceptibility to plasticity—rather than direct, such as through synapse formation or elimination. Regarding the first hypothesis, we could not find alternation of gene expression of astrocytic synapse organizers such as thrombospondins, Hevin, Chordin-like 1, or glypicans (Allen et al., 2012; Blanco-Suarez et al., 2018; Christopherson et al., 2005; Eroglu et al., 2009; Xu et al., 2010) in DEGs of hM3D vs. Ctrl or hM3D DR vs. LR. Rather, as shown in Figure 4, those regulating the ECM or adhesive functions, and affecting neuronal and synaptic functions, are included. For instance, *Igfbp2* (Caldwell et al., 2022; Shigetomi et al., 2024), *Igsf11* (Pelz et al., 2024), or *Dio2* (Bocco et al., 2016) in astrocytes have been reported to affect neuronal function. Therefore, the upregulation of these genes may contribute to the enhancement of RG synapses’ susceptibility to visual experience and to the reopening of the CP. Astrocytes are thought to contribute to the mechanisms underlying CP closure by stabilizing neural circuits through ECM maturation and cell-adhesion–related pathways (Ackerman et al., 2021; Ribot et al., 2021). It is therefore highly intriguing that genes related to the modulation of the extracellular environment were found among those upregulated by DREADD stimulation regardless of visual experience (Figure 4). Although DREADD stimulation does not revert astrocytes to a juvenile state, these findings suggest that it may enhance plasticity by modulating molecular pathways involved in CP closure. Notably, genes related to ECM regulation, such as Pcsk5 (Ito et al., 2021), differed from those implicated in CP closure mechanisms in the visual cortex. One possible explanation for this difference is the structural characteristic of the dLGN, which lacks perineuronal nets (PNNs) (Sabbagh et al., 2018) that plays a crucial role in the regulation of neuronal plasticity in the visual cortex (Pizzorusso et al., 2002) and other brain areas (Li et al., 2024; Sorg et al., 2016). If astrocyte stimulation acts on the perisynaptic environment or structure of RG synapses to regulate the permissiveness of plasticity, rather than astrocytes actively driving synapse formation or pruning themselves, it is likely that factors outside astrocytes—such as the presence or absence of neuronal activity associated with visual experience—play a dominant role in determining whether plasticity occurs and, if it does, the direction of synaptic remodeling.

We previously reported that in the primary somatosensory (S1) cortex of neuropathic pain model mice, astrocytes increased their Ca^2+^ activity and promoted synapse formation that underlie the emergence of chronic pain by secreting thrombospondin 1 through metabotropic glutamate receptor subtype 5 (mGluR5) activation (Kim et al., 2016). Using the same model, we also reported that, in the chronic phase of neuropathic pain, chemogenetic activation of astrocytes in the S1 cortex, combined with suppression of peripheral nerve activity by tetrodotoxin treatment, promoted elimination of synapses formed after peripheral nerve injury and cured allodynia (Takeda et al., 2022). These findings strongly suggest an astrocyte contribution to bidirectional synapse turnover under conditions requiring remodeling of neuronal circuits, while the underlying mechanisms remain unclear.

The potential of astrocyte stimulation to regain bidirectional neuronal plasticity that we reported here may enable control of neuronal circuit plasticity not only in sensory circuits during development but also in higher-order brain regions and pathological conditions. To translate the CP reopening via astrocyte stimulation into other experimental systems and clinical applications, it is necessary to clarify how these genes contribute to bidirectional synaptic reorganization, an area that remains a future challenge.

## Experimental procedures

### Animals

We used C57BL/6 mice (Japan SLC Inc.), Ai32 mice (JAX 012569, B6;129S-Gt(ROSA)26Sor^tm32(CAG-COP4*H134R/EYFP)Hze^/J) (Madisen et al., 2012) and Chx10 BAC mice (JAX 005105, STOCK Tg(Chx10-EGFP/cre,-ALPP)2Clc/J) (Rowan and Cepko, 2004). To drive the expression of ChR2 in RGCs, homozygous or heterozygous Chx10 BAC mice were crossed with homozygous Ai32 mice as previously described (Litvina and Chen, 2017). Their progeny heterozygous for each gene expressing ChR2 in RGCs (Chx10-cre; ChR2) were used for the experiments. Mice were group-housed with their litters on a 12hr light/dark cycle with ad libitum access to standard rodent chow and water before experiments. During light-rearing (LR) or dark-rearing (DR), mice were transferred to a box with an independent lighting system. For light-rearing group, mice were raised in standard conditions under a 12-hr light/dark cycle. For dark-reared mice, the light was turned off for 24 hrs. All experiments were approved by the Animal Care and Use Committee of the National Institutes of Natural Sciences and performed according to institutional guidelines.

### Generation of Adeno-Associated Viruses (AAVs)

Custom AAV vectors were constructed to enable selective expression of chemogenetic and fluorescent reporters in astrocytes. The viral plasmid pAAV-hALDH1L1-hM3D(Gq)-P2A-T2A-mScarlet-WPRE3 was generated to express the Gq protein-coupled DREADD receptor hM3D (Armbruster et al., 2007) and the red fluorescent protein mScarlet under the control of the human ALDH1L1 promoter (hALDH1L1), which ensures astrocyte-specific expression (Koh et al., 2017). The construct was packaged into a capsid of the engineered serotype AAVF (Beharry et al., 2022), a mutant derivative of AAV9, to enhance astrocyte targeting. A negative control vector, AAVF-hALDH1L1-mScarlet-WPRE3, expressing only mScarlet under the same promoter, was similarly constructed.

Plasmid production and AAV packaging were performed via triple transfection of HEK293T cells with the expression vector, an AAVF capsid helper plasmid, and adenoviral helper plasmids. Seventy-two hours post-transfection, viral particles were harvested from both the media and cell lysate, purified using iodixanol gradient ultracentrifugation, and concentrated using Amicon Ultra centrifugal filters (Millipore). Genomic titers were quantified via qPCR targeting the WPRE sequence. Final titers were determined to be 4.28 × 10⁹ genome copies (GC)/µL for AAVF-hALDH1L1-hM3D(Gq)-P2A-T2A-mScarlet-WPRE3 and 1 × 10¹⁰ GC/µL for AAVF-hALDH1L1-mScarlet-WPRE3. Viral aliquots were stored at −80°C until use.

## Method Details

### Immunohistochemistry

Chx10-cre; ChR2 mice were sacrificed two weeks after virus injection. Following perfusion, brains were dissected, fixed in 4% PFA at 4℃ overnight, then soaked sequentially in 30% sucrose overnight, and finally frozen in optimal cutting temperature compound (O.C.T., Sakura Finetek Japan). The coronal brain slices containing dLGN (16 µm in thickness) were prepared using a Leica CM3050s cryostat.

The brain slices were rehydrated in Phosphate Buffered Saline containing 0.1% Tween 20 (PBS-T). For immunofluorescence staining, the brain slices were blocked with 5% Bovine Derum Albumin (BSA; A9647, Sigma) dissolved in PBS-T. Primary antibody, rabbit anti-S100β antibody (marker for astrocytes, 1:100; ab52642, Abcam), was incubated with samples overnight at 4℃. After washing slices 3 times with PBS-T, samples were incubated with Goat anti-Rabbit IgG (H+L) Cross-Adsorbed Secondary Antibody, Alexa Fluor™ 405 (1:500; A-31556, Invitrogen) for 1 hr at room temperature (R.T.). To visualize neurons, the brain sections were incubated in NeuroTrace™ 435/455 Blue Fluorescent Nissl Stain (1:200; N21479, Invitrogen) for 20 min at R.T. after rehydration. Stained sections were mounted in VECTASHIED Hard Set Mounting medium (H-1400; Vector Laboratories). Confocal microscopy was carried out on Nikon A1R microscope equipped with 10x-20x objectives. For quantification, the cell numbers were counted manually with imageJ.

### Ca²^+^ Imaging of Astrocytes in Cultured Hippocampal Slices

Hippocampal slice cultures were prepared as described previously (Ueda et al., 2022). Briefly, hippocampal slices (350 µm thick) were obtained from postnatal day 6–9 C57BL/6N mice. The slices were cultured on membrane inserts (PICM0RG50; Millipore, Darmstadt, Germany) placed over culture medium containing 50% minimal essential medium (MEM), 21% Hank’s balanced salt solution (HBSS), and 25% horse serum, supplemented with (in mM): 15 NaHCO_3_, 6.25 HEPES, 10 D-glucose, 1 L-glutamine, 0.88 ascorbic acid, and 1 µg/mL insulin. Cultures were maintained at 35 °C in a humidified incubator with 5% CO_2_. Slices were transduced after 2–6 days *in vitro* with either AAVF-hALDH1L1-hM3D(Gq)-P2A-T2A-mScarlet-WPRE3 or AAVF-hALDH1L1-mScarlet-WPRE3, together with a Ca²^+^ indicator virus (AAVDJ-gfaABC1D-jGCaMP8m-WPRE3 produced by Murakoshi’ lab). Viral delivery was performed using a glass micropipette (Narishige, Tokyo, Japan).

Ca²⁺ imaging was performed under an epifluorescence microscope. For excitation, a blue light (475 nm LED; CoolLED, UK) for jGCaMP8m, or green light (565 nm LED; CoolLED, UK) for mScarlet was used. Time-lapse images were acquired using a sCMOS camera (ZYLA 4.2; Andor, UK) mounted on a microscope (BX51WI; Evident co., Japan) through a 20x water immersion objective lens at a rate of 2 frames/sec. Slices were continuously perfused with an imaging buffer solution (136 mM NaCl, 5 mM KCl, 0.8 mM KH_2_PO_4_, 20 mM NaHCO_3_, 1.3 mM L-glutamine, 0.2 mM ascorbic acid, 2 mM CaCl_2_, 2 mM MgCl_2_, MEM amino acids solution [Gibco; Thermo Fisher, USA], MEM vitamin solution [Gibco; Thermo Fisher, USA], and 1.5 mg/mL phenol red) aerated with 95% O_2_/5% CO_2_ at 24–26°C. A 30-s baseline recording was obtained before application of 10 μM clozapine-N-oxide (CNO), which was perfused into the recording chamber. Imaging continued for an additional 2.5 min after CNO application. Regions of interest (ROIs) were drawn around astrocytes co-expressing mScarlet and jGCaMP8m. Fluorescence intensity was quantified using ImageJ software. Changes in intracellular Ca²^+^ were expressed as percentage change relative to baseline fluorescence (F**_0_**) according to the formula:

ΔF/F**_0_** (%) = [(F – F**_0_**)/F**_0_**] × 100.

Only ROIs showing robust co-localization of the reporter and the indicator were included in the analysis.

### Stereotaxic viral injection

Chx10-cre; ChR2 mice at their age of P14 were anesthetized with 1.5-1.8% isoflurane (Viatris Inc., Tokyo, Japan) and placed in a stereotaxic frame (Narishige, Japan). DREADD virus (AAVF-hALDH1L1-hM3D(Gq)-P2AT2A-mScarlet-WPRE3, titer: 4.28×10^9^ GC/µL) or control virus (AAVF- hALDH1L1 -mScarlet-WPRE3, titer: 1×10^10^ GC/µL) diluted in saline were injected using NanoFil 34-gauge beveled needle (WPI, Sarasota, FL, USA). The virus-containing solution (200 nl) was injected to the dLGN (ML ±2.2 mm, AP +1.4 mm, DV -2.6 mm from the Lambda) at a rate of 100 nl/min with a stereotaxic injector (QSI; Stoelting, Wood Dale, IL, USA). At the end of the injection, the needle remained in place for 5 min to ensure the viral diffusion.

### CNO treatment

Clozapine N-oxide (CNO; 1 mg/kg, diluted in saline) was administered intraperitoneally (i.p.) to mice at 12-hour intervals. For *in vivo* administration, an infrared night scope was used when we performed CNO or saline administration to mice during dark rearing. For Ca^2+^ imaging with cultured hippocampal slices, 10 µM CNO was applied to the perfusion solution.

### Slice preparations and Whole-cell recordings

For slice preparation, Chx10-Cre; ChR2 mice aged postnatal day (P) 29–46 were anesthetized via intraperitoneal (i.p.) injection with a mixed anesthetic solution containing 40% ketamine hydrochloride (Ketalar® injection 500 mg, Daiichi Sankyo Co., Ltd., Tokyo, Japan) and 15% xylazine hydrochloride (Selactar® 2% injection, Elanco Japan K.K., Tokyo, Japan), diluted in saline. Mice were sacrificed after one week of either dark rearing or light rearing, depending on the experimental condition. Animals were intracardially perfused with ice-cold cutting solution containing (in mM): 93 NMDG, 2.5 KCl, 1.2 NaH_2_PO_4_, 30 NaHCO_3_, 20 HEPES, 25 glucose, 5 sodium ascorbate, 3 sodium pyruvate, 0.5 CaCl2, 10 MgSO4, and 12 N-acetyl-L-cysteine (pH 7.3), bubbled with 95% O_2_ and 5% CO_2_. Parasagittal brain slices (250–300 µm thick) containing the dorsal lateral geniculate nucleus (dLGN) were prepared using a Neo LinearSlicer MT (D.S.K). The slices were transferred to warmed (30 °C) oxygenated artificial cerebrospinal fluid (ASCF) containing (in mM): 125 NaCl, 2.5 KCl, 1.25 NaH_2_PO_4_, 26 NaHCO_3_, 20 glucose, 1 MgSO_4_, and 2 CaCl_2_ for 30 min for recovery and subsequently maintained at room temperature until use.

Whole-cell voltage-clamp recordings were performed from thalamocortical (TC) neurons using a Multiclamp 700B amplifier and digitized with a Digidata 1440A interface (Molecular Devices, San Jose, CA, USA). Signals were sampled at 20 kHz and acquired with pClamp 11.2 software (Molecular Devices). Recordings were conducted at room temperature using a Nikon E600FN microscope (Nikon Instruments Inc., Melville, NY, USA) equipped with an IR-CCD camera (IR1000, DAGE-MTI, Michigan City, IN, USA) and an LED light source. Slices were continuously perfused with oxygenated ACSF containing pharmacological agents to isolate excitatory postsynaptic currents (EPSCs). To pharmacologically isolate EPSCs, the following antagonists and modulators were bath-applied: 20 µM bicuculline methochloride (GABA_A_ receptor antagonist; Tocris, Cat. No. 0131), 50 µM cyclothiazide (AMPAR desensitization blocker; Tocris, Cat. No. 0713), 10 µM (R)-CPP (NMDA receptor antagonist; Tocris, Cat. No. 0247), 10 µM DPCPX (adenosine A_1_ receptor antagonist; Tocris, Cat. No. 0439), 1 µM LY341495 (mGluR2/3 antagonist; Tocris, Cat. No. 1209), and 1 µM CGP55845 (GABA_B_ receptor antagonist; Tocris, Cat. No. 1248). All compounds were prepared as stock solutions according to the manufacturer’s instructions and diluted to their final concentrations in ACSF immediately before use.

Patch electrodes (2–4 MΩ) were filled with a internal solution containing (in mM): 35 CsF, 95 CsCl, 10 EGTA, 10 HEPES, 4 ATP, 0.4 GTP, 5 lidocaine, and 0.1 methoxyverapamil hydrochloride (pH 7.4, adjusted with CsOH, 285 mOsm). RG-EPSCs were evoked by optical stimulation and recorded at a holding potential of –30 mV. Full-field optical stimulation to evoke ChR2-mediated EPSCs was delivered using 470-nm light through a 40x objective. Light intensity was incrementally adjusted from 0.52 to 0.62 mW/mm² (1%) to the maximum intensity of 59.2 mW/mm² (100%) using an LED control panel.

### Single-nucleus RNA Sequencing (snRNA-seq)

For each experimental group, the dorsal lateral geniculate nucleus (dLGN) was microdissected from 4–6 mice and pooled to ensure sufficient nuclei yield. Nuclei were isolated using the Minute™ Detergent-Free Nuclei Isolation Kit for Animal Tissues (Invent Biotechnologies, Cat. No. NI-024, 20 reactions), following the manufacturer’s protocol to preserve nuclear integrity. The isolated nuclei were filtered and sorted using a Sony SH800 Cell Sorter to enrich for single, intact nuclei prior to library preparation.

Single-nucleus RNA-seq libraries were prepared using the Chromium Next GEM Single Cell 3′ HT Reagent Kits v3.1 (10x Genomics, Cat. No. 1000370), along with the Chromium Next GEM Chip M Single Cell Kit (Cat. No. 1000371) and Dual Index Kit TT Set A (Cat. No. 1000215), according to the manufacturer’s instructions. Quality control of amplified cDNA and final libraries was performed using an Agilent Bioanalyzer High Sensitivity DNA Kit and Qubit dsDNA High Sensitivity Assay Kit.

Sequencing was conducted by Azenta Life Sciences (New York City, NY) on an Illumina platform. Raw sequencing data were processed using the Cell Ranger pipeline (10x Genomics) for alignment, barcode assignment, and UMI counting. Downstream analysis, including normalization, clustering, and cell type identification, differential expression (DE) testing was performed using the Seurat package (version 5.2.1) in R.

We performed over-representation analysis (ORA) to assess the functional enrichment of Gene Ontology (GO) biological process (BP) gene sets among the identified differentially expressed genes (DEGs). Enrichment was carried out using the enrichGO function from the clusterProfiler package (Wu et al., 2021; Yu et al., 2012) with the following parameters: OrgDb = org.Mm.eg.db, ont = "BP", pAdjustMethod = "BH", pvalueCutoff = 0.05, and qvalueCutoff = 0.05.

Given the large number and redundancy of enriched GO terms, we used rrvgo to cluster ORA results (Sayols, 2023), following a modified version of a published method (Nelson et al., 2023). Semantic similarity was computed using the Wang method (Wang et al., 2007), and biological process terms were clustered at a similarity threshold of 0.9. Within each cluster, the term with the highest - log_10_(adj. p-value) was designated as the parent term. Terms in each cluster were subsequently ranked by -log_10_(adj. p-value), and top terms were selected to minimize redundancy in meaning or functional role.

Functional analysis of the common gene set was performed using Metascape (https://metascape.org/). In the Membership section, we searched for the keyword “adhesion” within the term names and descriptions across multiple annotation sources, including GO Biological Process, Reactome Gene Sets, KEGG Pathway, WikiPathways, and PANTHER Pathway, to identify genes associated with adhesion-related biological functions.

## Quantification and Statistical Analysis

For electrophysiological data, statistical analyses and data visualization were performed in Excel. Statistical comparisons between two groups were performed using the Mann–Whitney U-test. Data are presented as mean ± SEM. Statistical significance was defined as p < 0.05.

## Data availability

The RNA-seq datasets generated in this study will be made publicly available in the Gene Expression Omnibus (GEO) after acceptance of the peer-reviewed manuscript.

## Acknowledgement

This work was supported by JSPS Grants-in-Aid for Scientific Research (KAKENHI) grant numbers: 20H05916 and 24K09541(M.N.), 21H05245 (Y.G.), 20H00500 and 24K02213 (J.N.); The Naito Foundation (M.N.). We thank Dr. Hideji Murakoshi and Dr. Hiromi Ueda, Mr. Yutaro Nagasawa, and Ms. Tatsuko Ohba (NIPS) for helping with AAV production and Ca^2+^ imaging (supported by JSPS KAKENHI Grant Number JP22H04926), Drs. Hirokazu Hirai and Yasunori Matsuzaki (Gumma University) for producing AAVs with hALDH1L1 promoter (supported by AMED Grant Number JP21dm0207111), Ms. Kyoko Noguchi, Ms. Yuki Ichihashi, and Ms. Junko Ishida (NIPS) for snRNA-seq technical assistance, and Dr. Hiroaki Wake (Nagoya University and NIPS) and Dr. Mariko Miyata (Tokyo Women’s Medical University) for helpful comments.

## Author contributions

C.J., J.N., and M.N. conceived and designed the project. C.J., M.N., and Y.G. performed the experiments and analyzed the data. C.J., Y.G., and M.N. wrote the manuscript. All the coauthors discussed the results and exchanged comments on the manuscript.

## Declaration of interests

The authors declare no competing interests.

## Notes

### Competing Interest Statement

The authors have declared no competing interest.

