## Supplementary material for "Astrocyte stimulation reopens the window of the critical period for the experience-dependent plasticity of retinogeniculate synapses": KRT69250cf247_JIN et al_20260111.docx

### Key resources table

| REAGENT or RESOURCE | SOURCE | IDENTIFIER |
| --- | --- | --- |
| Antibodies | | |
| Rabbit anti-S100β | Abcam | RRID:AB_882426 |
| Goat anti-Rabbit IgG (H+L) Cross-Adsorbed Secondary Antibody, Alexa Fluor™ 405 | Invitrogen | RRID:AB_221605 |
| NeuroTrace™ 435/455 Blue Fluorescent Nissl Stain | Invitrogen | Cat# N21479 |
| Bacterial and virus strains | | |
| AAVF-hALDH1L1-hM3D(Gq)-P2AT2A-mScarlet-WPRE3 | This paper | N/A |
| AAVF-hALDH1L1 -mScarlet-WPRE3 | This paper | N/A |
| AAVDJ-gfaABC1D-jGCaMP8m-WPRE3 | Section of Multiphoton Neuroimaging, National Institute for Physiological Sciences | N/A |
| Chemicals, peptides, and recombinant proteins | | |
| (-)-Bicuculline methochloride | Tocris | Cat# 0131 |
| Cyclothiazide | Tocris | Cat# 0713 |
| (*R*)-CPP | Tocris | Cat# 0247 |
| DPCPX | Tocris | Cat# 0439 |
| LY 341495 | Tocris | Cat# 1209 |
| CGP 55845 hydrochloride | Tocris | Cat# 1248 |
| Clozapine *N*-oxide (CNO) | Tocris | Cat# 4936 |
| Bovine Serum Albumin | Sigma Aldrich | Cat# A9647 |
| Critical commercial assays | | |
| Chromium Next GEM Single Cell 3’HT Reagent Kits v3.1 | 10X Genomics | Cat#1000370 |
| Chromium Next GEM Chip M Single Cell Kit | 10X Genomics | Cat#1000371 |
| Bioanalyzer High Sensitivity DNA Kit | Agilent | Cat#5067-4627 |
| Qubit™ dsDNA High Sensitivity Assay Kit | Thermo Fisher Scientific | Cat#Q32854 |
| Agilent 2100 Bioanalyzer | Agilent | Cat#G2938C |
| Qubit® 2.0 Fluorometer | Thermo Fisher Scientific | Cat#Q32866 |
| Deposited data | | |
| Raw snRNA-seq data | This paper | GEO: GSE316066 |
| Experimental models: Organisms/strains | | |
| Mouse: C57BL/6J | Japan SLC Inc. | N/A |
| Mouse: B6;129S-Gt(ROSA)26Sor^tm32(CAG-COP4*H134R/EYFP)Hze^/J | The Jackson Laboratory | RRID:IMSR_JAX:012569 |
| Mouse: STOCK Tg(Chx10-EGFP/cre,-ALPP)2Clc/J | The Jackson Laboratory | RRID:IMSR_JAX:005105 |
| Software and algorithms | | |
| pClamp v11.2 | Molecular Devices | RRID:SCR_011323 |
| FIJI | NIH ImageJ software | http://fiji.sc/ |
| Excel | Microsoft | N/A |
| Cell Ranger v9.0.0 | 10x Genomics | https://www.10xgenomics.com/support/software/cell-ranger/latest |
| R version v4.5.1 | The R project | https://cran.r-project.org/ |
| RStudio v2025.09.2+418 | R Studio, PBC | RStudio Desktop - Posit |
| Seurat v5.3.1 | Hao et al. 2023 | https://satijalab.org/seurat/ |
| clusterProfiler v4.14.6 | Wu et al., 2021 | https://bioconductor.org/packages/release/bioc/html/clusterProfiler.html |
| rrvgo v1.18.0 | Sayols, 2023 | https://www.bioconductor.org/packages/release/bioc/html/rrvgo.html |
| Other | | |
| Multiclamp 700B amplifier | Molecular Devices | N/A |
| Digidata 1440A interface | Molecular Devices | N/A |
| Nikon E600FN microscope | Nikon Instruments Inc | N/A |
| IR-CCD camera | DAGE-MTI | IR1000 |
| Sony SH800 Cell Sorter | Sony Biotechnology | RRID:SCR_018066 |
