## Supplementary material for "Astrocyte stimulation reopens the window of the critical period for the experience-dependent plasticity of retinogeniculate synapses": Supplemental information_Jin et al_20260111.docx

**Supplemental Figures:**

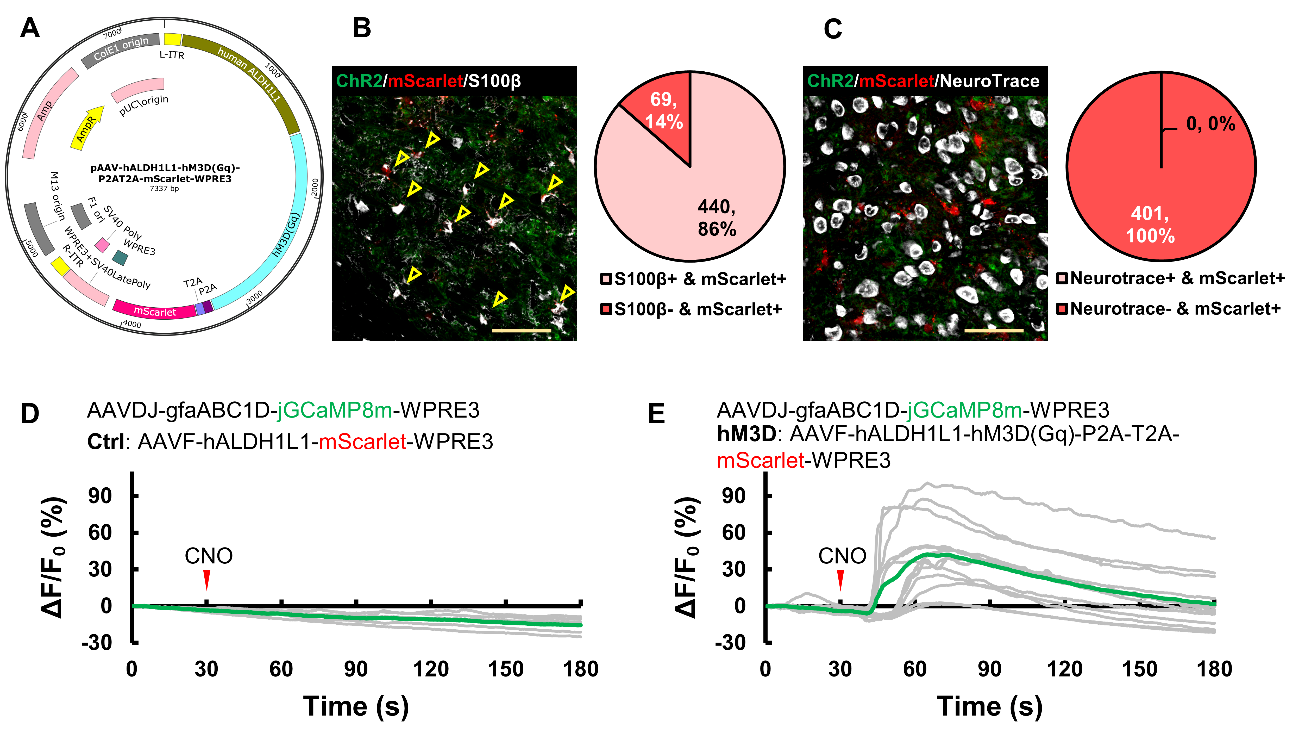

**Figure S1 (related to Figures 2 and 3). Development and validation of Gq virus expression in thalamic astrocytes.**

**(A)** A scheme of the viral construct. The vector expresses hM3D under the human ALDH1L1 (hALDH1L1) promoter, with mScarlet as a marker protein linked via P2A-T2A. The control construct lacked the hM3D and P2A-T2A sequences, expressing only mScarlet.

**(B–C)** Expression of mScarlet (red) in the dLGN. Two weeks after viral infection in Chx10-Cre;ChR2 mice, brain slices containing the dLGN were prepared following fixation with 4% paraformaldehyde. The green signal indicates ChR2–EYFP expression in retinal fibers. Astrocytes (B) and neurons (C) were labeled with S100β antibody (B, left panels, white) or NeuroTrace (C, left panels, white). Pie charts (right) show the percentage of S100β-positive (B) or NeuroTrace-positive (C) cells among mScarlet-positive cells. Overall, 86% of mScarlet-positive cells were S100β-positive, whereas no NeuroTrace-positive cells expressed mScarlet, indicating that most mScarlet-expressing cells were astrocytes driven by the hALDH1L1 promoter. Quantification was performed on 3–5 mice, using at least two mScarlet-expressing dLGN sections per mouse. Scale bars, 50 μm.

**(D-E)** Confirmation of Ca²⁺ elevation with CNO application in cultured hippocampal slices infected with hALDH1L1 hM3D mScarlet or hALDH1L1 mScarlet AAVs. AAV-gfaABC1D-jGCaMP8m-WPRE3 was used to monitor Ca²⁺ elevation in astrocytes. CNO was applied after 30 seconds of baseline recording, and changes in fluorescence intensity were analyzed. Green or grey lines represent Ca²⁺ responses of averaged (green) or each cell (grey) recorded from 13 (hM3D) or 6 (Ctrl) cells. CNO application induced Ca²⁺ responses successfully in hM3D mScarlet-expressing cultured cells.

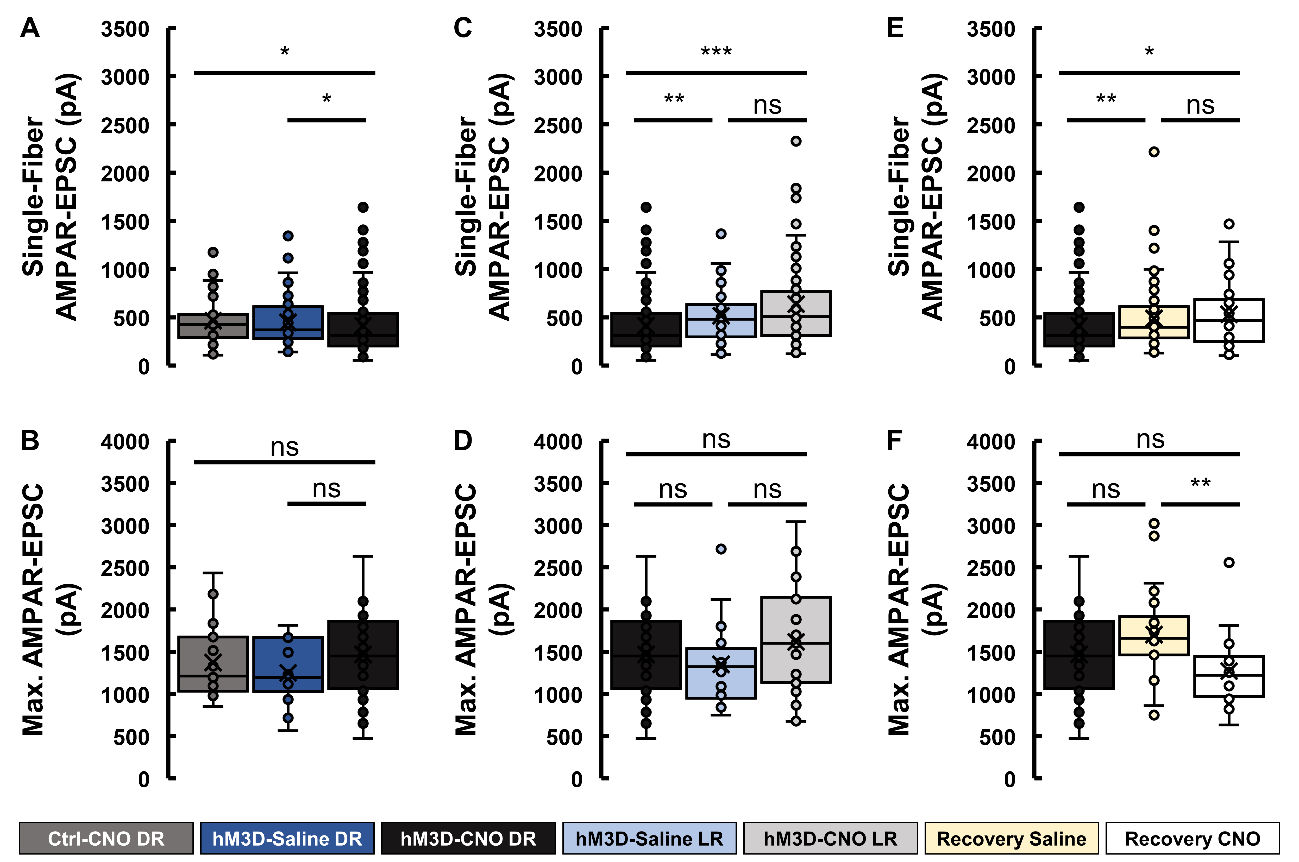

**Figure S2 (related to Figures 2 and 3). Properties of retinogeniculate EPSCs in Chx10-Cre;ChR2 mice in post–critical period.**

Single-fiber AMPAR-mediated EPSC amplitude (A, C, E) and maximal AMPAR-mediated EPSC amplitude (B, C,F) recorded at a holding potential of –30 mV. All boxplots represent the distribution of EPSC amplitudes according to the following conventions:

The horizontal line inside each box indicates the median value. The upper and lower edges of the box represent the first and third quartiles. Whiskers extend to the most extreme data points within 1.5× the interquartile range. Data points plotted outside the whiskers represent outliers. Mean values are shown as “×” marks. Individual data points are displayed as open circles. (A and D) Boxplots representing single-fiber EPSC amplitudes (A) and maximum amplitudes (D) in Ctrl-CNO DR (grey), hM3D-saline DR (blue), and hM3D-CNO DR (black) conditions. (B and E) The same analysis as (A) and (D) in hM3D-CNO DR (black), hM3D-saline LR (pale blue), and hM3D-CNO LR (pale grey) conditions. (C and F) The same analysis as (A) and (D) in hM3D-CNO DR (black), Recovery saline (pale yellow) and Recovery CNO (white) conditions. *, **, or *** represent p < 0.05, 0.01, or 0.001 with Mann-Whitney U-test; ns, not significant.

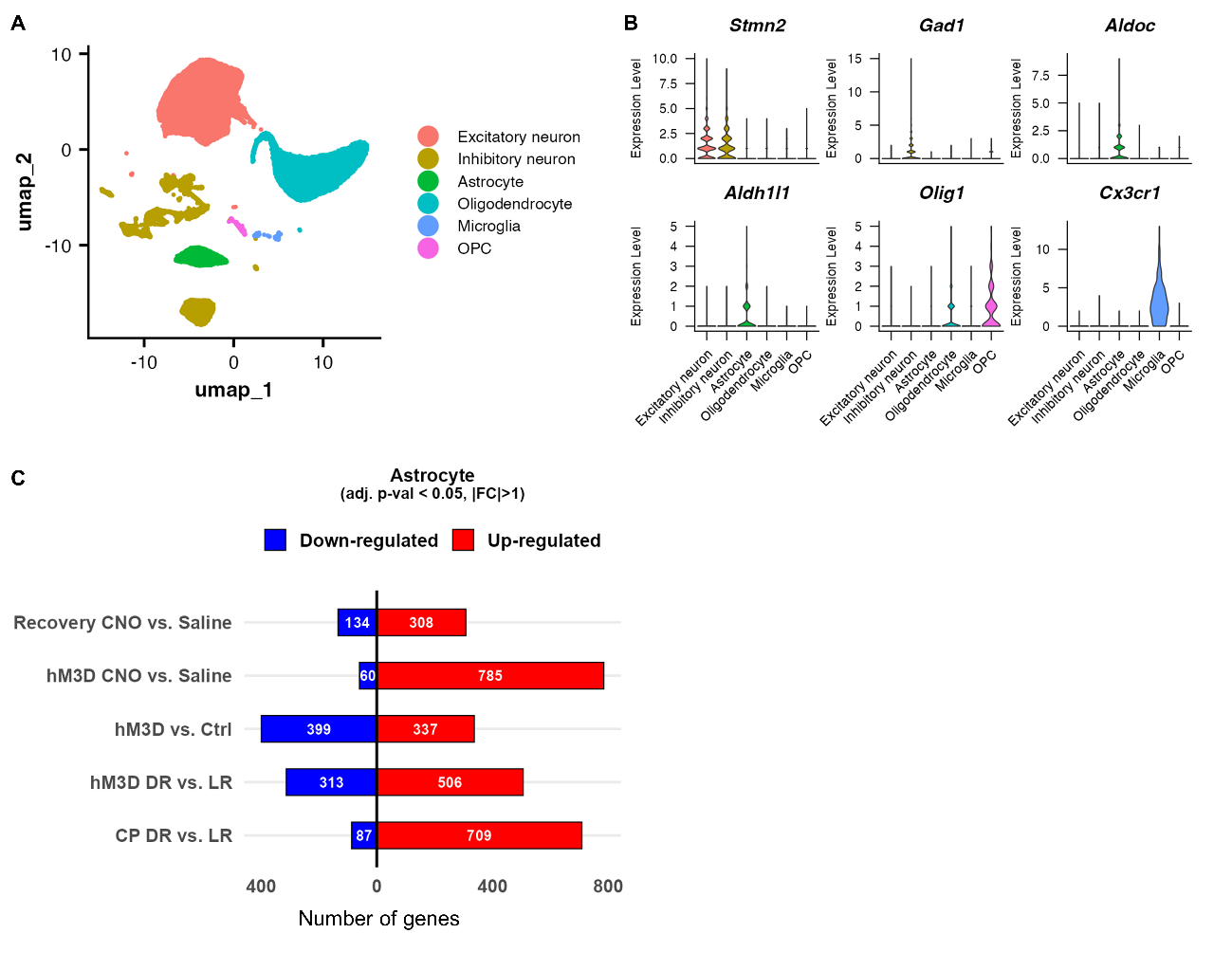

**Figure S3 (related to Figure 4). Classification of cell types and differential expression analysis**

**(A)** UMAP visualization of all cells, colored by annotated cell type. Major populations include excitatory neurons, inhibitory neurons, astrocytes, oligodendrocytes, and oligodendrocyte progenitor cells (OPC). Cell type annotation was based on canonical marker gene expression and clustering results.

**(B)** Violin plots showing expression levels of representative marker genes used for cell type identification: *Stmn2* for excitatory neurons, *Gad1* for inhibitory neurons, *Aldoc* and *Aldh1l1* for astrocytes, *Olig1* for oligodendrocytes and OPC, and *Cx3cr1* for microglia.

**(C–D)** Diverging bar plots showing the numbers of up- and down-regulated differentially expressed genes (DEGs) for the corresponding comparisons in dLGN astrocytes. DEGs were defined as those with an adjusted p-value < 0.05 and an absolute fold change (FC) > 1.

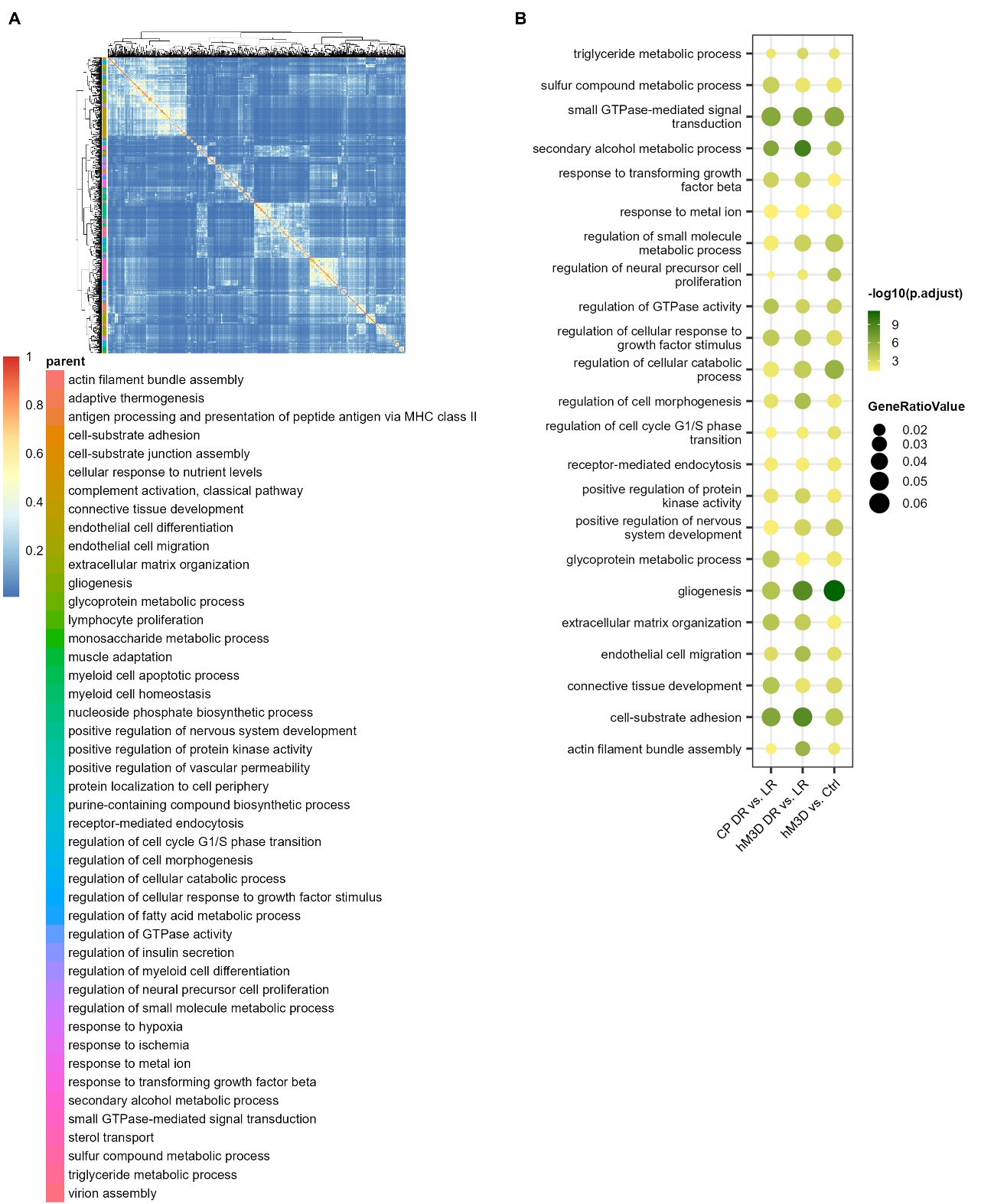

**Figure S4 (related to Figures 4). GO over-representation analysis (ORA) results and clustering for astrocytes**

(A) Heatmap showing the similarity matrix generated from semantic similarity clustering analysis of all statistically significant BP GO terms, pooled across the astrocytes for the CP DR vs. LR, hM3D DR vs. LR, and hM3D vs. Ctrl comparisons. Semantic similarity analysis (*rrvgo*) was used to reduce redundancy among enriched GO terms by grouping them according to their relatedness within the GO graph and defining representative “parent terms” for each cluster. The selected BP terms were defined as parent terms, and they were used to analyze –log10(adjusted p-value) and the gene ratio shown in (B).

(B) Dot plots showing the common parent terms among these three comparison pairs based on semantic similarity analysis of ORA results. Circle color represents –log10(adjusted p-value), and circle size corresponds to the gene ratio associated with each term.

**Supplemental Tables:**

| **Related to** | **Animal** | **Age** | **# Cell** | **Single-Fiber AMPAR-EPSC (pA)** | **Max. AMPAR-EPSC (pA)** | **Single-Fiber  Fraction** | **PPR** |
| --- | --- | --- | --- | --- | --- | --- | --- |
| Figure 1 | CP LR | P29-38 | 20 | 536.77 ± 42.95 | 1610.31 ± 144.91 | 0.33 ± 0.02 | 0.62 ± 0.02 |
|  | CP DR | P29-36 | 27 | 337.65 ± 16.64 | 1363.11 ± 79.65 | 0.25 ± 0.01 | 0.63 ± 0.01 |
|  | post-CP LR | P36-43 | 15 | 529.53 ± 52.72 | 1341.48 ± 125.84 | 0.39 ± 0.04 | 0.69 ± 0.02 |
|  | post-CP DR | P35-42 | 16 | 495.98 ± 47.95 | 1146.96 ± 77.16 | 0.43 ± 0.03 | 0.67 ± 0.02 |
| Figure 2 | Ctrl-CNO DR | P36-42 | 19 | 465.48 ± 35.03 | 1371.93 ± 99.90 | 0.34 ± 0.02 | 0.61 ± 0.02 |
|  | hM3D-Saline DR | P36-46 | 23 | 448.64 ± 30.56 | 1248.40 ± 72.86 | 0.36 ± 0.02 | 0.61 ± 0.02 |
|  | hM3D-CNO DR | P35-41 | 35 | 414.10 ± 27.47 | 1467.11 ± 87.26 | 0.28 ± 0.02 | 0.66 ± 0.01 |
|  | hM3D-CNO LR | P37-43 | 28 | 637.51 ± 56.10 | 1616.54 ± 119.27 | 0.39 ± 0.03 | 0.64 ± 0.01 |
|  | hM3D-Saline LR | P35-39 | 21 | 516.16 ± 36.79 | 1351.86 ± 104.70 | 0.39 ± 0.03 | 0.57 ± 0.02 |
| Figure 3 | Recovery Saline | P37-42 | 27 | 489.22 ± 31.96 | 1703.22 ± 98.60 | 0.29 ± 0.02 | 0.67 ± 0.01 |
|  | Recovery CNO | P37-41 | 18 | 531.41 ± 48.98 | 1269.50 ± 103.35 | 0.42 ± 0.04 | 0.65 ± 0.02 |

**Table S1 (related to Figures 1–3).** The age of the mice, number of cells, and mean values for electrophysiological properties of AMPAR-mediated RG-EPSCs recorded in the indicated conditions in Chx10-Cre;ChR2 mice.

| **Cell type** | **CP LR** | **CP DR** | **Recovery CNO** | **Recovery Saline** | **Ctrl-CNO DR** | **hM3D-Saline DR** | **hM3D-CNO LR** | **hM3D-CNO DR** | **Total** |
| --- | --- | --- | --- | --- | --- | --- | --- | --- | --- |
| Oligodendrocyte | 374 | 3252 | 2537 | 2689 | 1623 | 2573 | 2625 | 2009 | 17682 |
| Excitatory neuron | 519 | 5197 | 3628 | 4644 | 3711 | 4606 | 4915 | 2866 | 30086 |
| Astrocyte | 282 | 576 | 793 | 355 | 797 | 228 | 912 | 560 | 4503 |
| Inhibitory neuron | 103 | 1256 | 870 | 1134 | 753 | 987 | 1085 | 706 | 6894 |
| NA | 54 | 386 | 100 | 102 | 75 | 89 | 155 | 102 | 1063 |
| OPC | 132 | 99 | 21 | 28 | 23 | 33 | 51 | 44 | 431 |
| Microglia | 78 | 48 | 29 | 36 | 14 | 56 | 40 | 18 | 319 |
| Endothelial | 2 | 64 | 17 | 18 | 23 | 31 | 33 | 12 | 200 |
| Pericyte | 5 | 19 | 8 | 18 | 11 | 16 | 16 | 20 | 113 |

**Table S2. The number of nuclei identified with marker gene expression in indicated experimental conditions.** We used *Stmn2* for excitatory neurons, *Gad1* for inhibitory neurons, *Aldoc* and *Aldh1l1* for astrocytes, *Olig1* for oligodendrocytes, *Cx3cr1* for microglia, *Pdgfra* for OPC, *Cldn5* for endothelial cells, and *Vtn* for Pericytes. Nuclei that did not express any marker genes were defined as NA.

**Supplemental data 1 (related to Figure 4).** Combined ORA results for astrocyte comparison pairs as described in Figure 4A.

**Supplemental data 2 (related to Figure 4).** Semantic similarity clustering of ORA results for astrocytes. Data are provided as an .csv file. Each sheet contains clustering results for all enriched GO Biological Process (BP) terms.

**Supplemental data 3 (related to Figure 4).** Metascape analysis of genes related to the “adhesion” membership.

This Excel workbook contains terms associated with “adhesion” in Biological Process and Pathway databases, including Reactome, WikiPathways, Panther Pathways, and KEGG. Selected genes (membership = 1) are listed.
